# NOJAH: Not Just Another Heatmap for Genome-Wide Cluster Analysis

**DOI:** 10.1101/415398

**Authors:** Manali Rupji, Bhakti Dwivedi, Jeanne Kowalski

**Author notes:** Corresponding author (JK).

## Abstract

Since their inception, several tools have been developed for cluster analysis and heatmap construction. The application of such tools to the number and types of genome-wide data available from next generation sequencing (NGS) technologies requires the adaptation of statistical concepts, such as in defining a most variable gene set, and more intricate cluster analyses method to address multiple omic data types. Additionally, the growing number of publicly available datasets has created the desire to estimate the statistical significance of a gene signature derived from one dataset to similarly group samples based on another dataset. The currently available number of tools and their combined use for generating heatmaps, along with the several adaptations of statistical concepts for addressing the higher dimensionality of genome-wide NGS-derived data, has created a further challenge in the ability to replicate heatmap results. We introduce NOJAH (NOt Just Another Heatmap), an interactive tool that defines and implements a workflow for genome-wide cluster analysis and heatmap construction by creating and combining several tools into a single user interface. NOJAH includes several newly developed scripts for techniques that though frequently applied are not sufficiently documented to allow for replicability of results. These techniques include: defining a most variable gene set (a.k.a., ‘core genes’), estimating the statistical significance of a gene signature to separate samples into clusters, and performing a result merging integrated cluster analysis. With only a user uploaded dataset, NOJAH provides as output, among other things, the minimum documentation required for replicating heatmap results. Additionally, NOJAH contains five different existing R packages that are connected in the interface by their functionality as part of a defined workflow for genome-wide cluster analysis. The NOJAH application tool is available at http://bbisr.shinyapps.winship.emory.edu/NOJAH/ with corresponding source code available at https://github.com/bbisr-shinyapps/NOJAH/.

## Introduction

Data from next generation sequencing (NGS) technologies, such as whole genome sequencing, have created a level of dimensionality that has greatly exceeded that of prior, microarray-based genome-wide datasets, resulting in the need for innovative approaches to cluster analysis and heatmap construction. For this reason, several disparate methods have been developed to address such needs. For example, consensus clustering was introduced as a method for estimating the number of clusters [1, 2]. The concept of defining a ‘most variable’ gene set was introduced to address the much higher dimension of NGS data by filtering out genes with little to no differences among samples with respect to some molecular data type and performing a cluster analysis on the remaining, ‘core gene set.’ This approach has resulted in the use of several definitions applied to define a core gene set, most of which are insufficiently documented to enable their replicability. Other concepts in cluster analysis, such as silhouette widths for examining the tightness of clusters, though around for some time, have gained renewed interest for their use in defining a ‘core sample set’ within the context of genomic data cluster analysis, an approach that has been particularly useful when clustering many samples [3]. We have collectively placed these new approaches and new adaptions of existing methods for genome-wide cluster analysis and heatmap construction into the following general, genome-wide heatmap analysis workflow: 1) define a most variable gene set (a.k.a., ‘core genes’); 2) perform cluster analysis using core genes and construct heatmap of results; 3) estimate the number of clusters; 4) define a core sample set and update the heatmap using both core genes and core samples.

The ability to implement steps two through four of this workflow would require at a minimum, knowledge on how to download and separately run five preexisting R packages, not to mention knowing what to document from each to enable heatmap replicability and how to use each tool within the context of constructing a genome-wide heatmap. The first step for defining a set of most variable genes is by definition, variable, with no universally agreed upon meaning. In statistics alone, at least three measures of spread could be applied to define a most variable gene set: 1) variance (VAR), 2) median absolute deviation (MAD); and 3) inter-quartile range (IQR). Considering the lack of consensus on defining a ‘most variable’ gene set, a step that has become common in the literature, NOJAH offers the user an analysis approach to this task that is specific to the data and includes several options and visuals. For example, a genome-wide variant heatmap creates a challenge due to the general sparseness of the data, particularly if using somatic mutations. For this reason, we have created an integrated most variable analysis approach to help addresses the sparseness often associated with variant data to enable the defining of a reduced-dimension, most variable gene set in this case. In addition to the options available for the selection of a most variable gene set, there are also several options from which to choose when creating a heatmap such as distance choice, clustering method, and data scaling. As the number of options increases, it becomes ever more challenging to replicate results of heatmaps and as such, we have implemented as part of the standard NOJAH output pipeline a workflow that documents the options used in creating a heatmap.

In addition to a genome-wide cluster analysis workflow for a single data type, NOJAH includes an option for applying this workflow to several genome-wide data types from the same samples, such as RNA-Seq derived gene expression, methylation and copy number, and combines cluster results based on a cluster of clusters approach [3]. Since the interpretation from combining cluster analysis results is not the same as a single cluster analysis, we have also included in NOJAH several descriptive measures to help guide the meaning of the resulting cluster of clusters.

Once a heatmap is constructed with core genes and samples, one is often interested in the statistical significance of the gene set for which approaches vary and are loosely defined in the literature. We have included in NOJAH a bootstrap approach for estimating the statistical significance of a derived gene set in separating samples into groups as compared to random gene sets of the same size. NOJAH includes several other options such as the desire for more intricate heatmaps that display phenotype information other than that used for clustering to assess potential cluster associations in real time. Considering the increasing number of public repositories containing genome-wide data on an increasing number of samples, there is a greater demand for more intricate approaches to cluster analysis and heatmap construction. Thus, within the context of current ‘big data,’ we refer to a heatmap construction of cluster analysis results as a ‘heatmap analysis. To address these and other increased needs, we developed NOJAH as a comprehensive genome-wide *heatmap analysis* tool using a web interface, making it Not Just Another Heatmap.

## Method and Results

### Analysis Workflows

NOJAH is organized into three separate analysis workflows that correspond to the use cases highlighted in Fig 1: (1) genome-wide heatmap (GWH) analysis, (2) combined results cluster(CrC) analysis, and (3) gene set significance of cluster (SoC) analysis. To demonstrate each use case, we obtained data from two public domain sources: 1) TCGA breast cancer (BRCA) data portal and 2) Multiple Myeloma Research Foundation (MMRF) coMMpass trial. In this paper, we use the TCGA breast cancer dataset to demonstrate the application of NOJAH to address each use case in Fig 1 (see *Supplementary Section* for NOJAH applications to coMMpass trial data).

**Fig 1.**
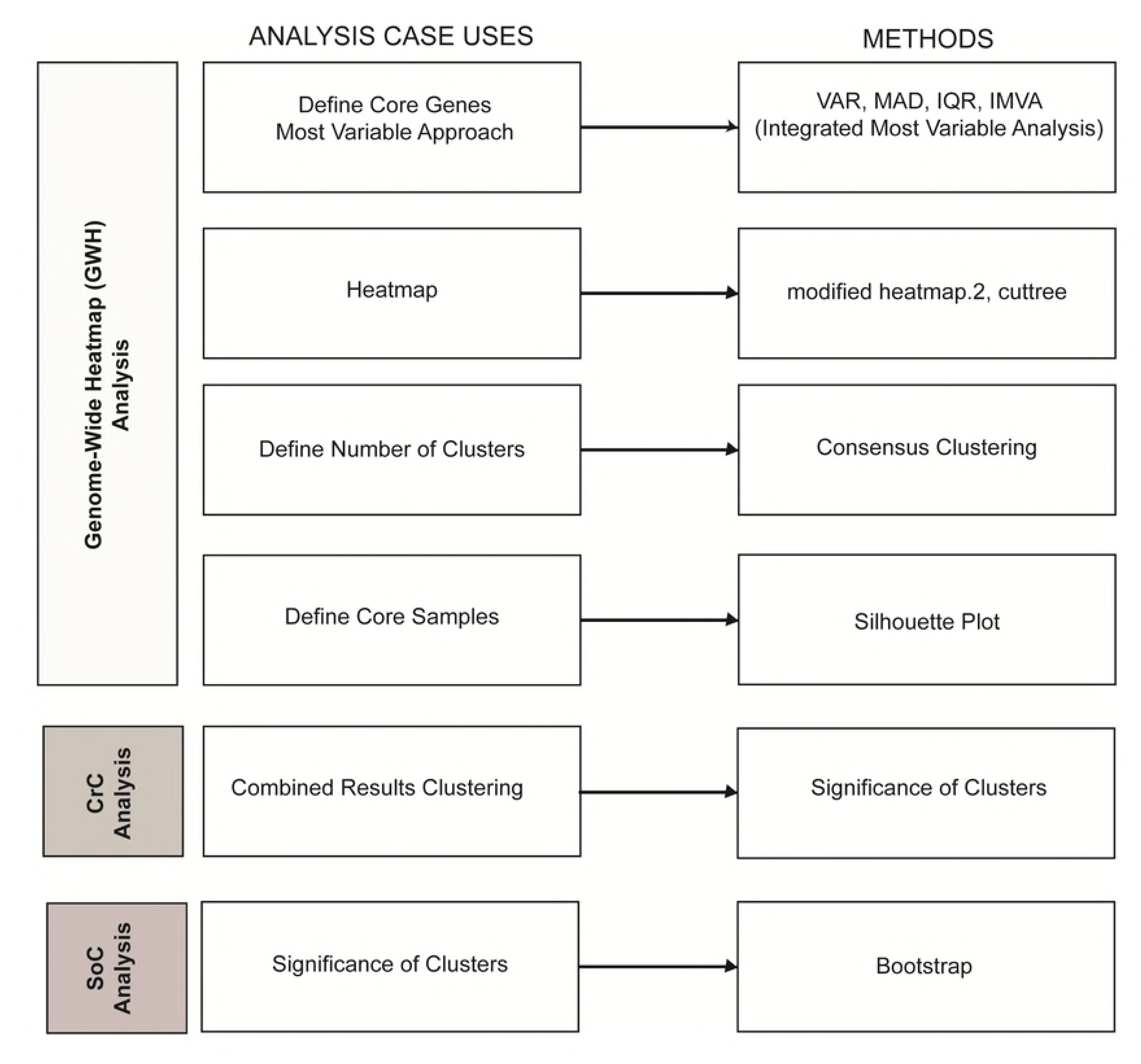
NOJAH Heatmap Analysis Use Cases. Workflows available in NOJAH to perform a genome-wide heatmap analysis of a single molecular data type (white tab), several molecular data types using a combined results cluster analysis (light grey), and for estimating the statistical significance of a derived gene set in separating sample groups (grey). The methods implemented in NOJAH to address each workflow are listed alongside each component case.

### Example TCGA BRCA Data

TCGA breast cancer RNA-Seq gene level expression data was downloaded from GDC data portal using TCGAbiolinks [4, 5]. Matched breast primary tumor-normal samples with available clinical, gene expression quantification data, 450K methylation and Copy Number Variation (CNV) information were extracted, resulting in total of 75 matched samples. Preprocessed FPKM RNA-Seq expression data was downloaded from the GDC data portal and was log2(count+1) transformed. Beta values from the 450K methylation data were downloaded and transformed using log2 ((1+beta)/(1-beta)). Copy segment mean values were downloaded from the GDC data portal and transformed by adding the lowest segment mean value among all samples to each copy segment.

Among the 75 breast cancer samples, 25 had an event of death reported. Among these 25 patients, a survival time of 6.2 years was identified as a cut point based on a martingale residual approach [6] for categorizing patients into those who died “early” (n=17) versus “late” (n=8). Patients were additionally classified into four different subtypes: basal-like (n=3), Her2 (n=5), LumA (n=5) and LumB (n=12) using PAM50 [7]. We illustrate the heatmap analysis workflows available in NOJAH, as illustrated in Fig 1 based on these 25 patients two sample groups.

### Genome-Wide Heatmap (GWH) Analysis

Genome-wide heatmaps are widely used to graphically display potential underlying patterns within the large genomic dataset. They have been used to reveal information about how the samples/genes cluster together and provide insights into potential sample biases or other artifacts. With genome-wide data, a heatmap analysis requires several steps to obtain a result. We first elaborate on each analysis case use defined within our GWH analysis workflow in Fig 1, followed by the application of each to the TCGA breast cancer data. First, a ‘topmost variable’ gene set is defined for characterizing differences with respect to some quantitative value among the sample groups. There is very poor to minimal documentation as to how these topmost variable subsets of genes are selected [3, 8-10] and yet, defining such a set is especially important as it is typically the first filter applied prior to conducting a cluster analysis. Though some tools provide a single measure of variability which aids in the filtering process, there are in fact, several measures of spread that could be applied and several ways to define cut points based on them for selecting a ‘topmost variable’ gene set. Some of the commonly used measures are variance (VAR) and median absolute deviation (MAD). These measures however will not work well for variant data which is critical to cancer research, since such data is sparse in nature, requiring further consideration for defining the notion of a variable gene set. Regardless of approach, a heatmap is then constructed based on a defined, ‘topmost variable’ subset of genes. Next, one often examines the number of clusters. Since genomic heatmaps are more commonly based on hierarchical clustering approaches, there is little to no confidence that the number of clusters estimated exists in the dataset and whether they depend on the choice of the clustering and distance measures. To examine the number of clusters in a dataset, consensus clustering is a popular method of choice. With the number of clusters estimated, the next question becomes one of how ‘tight’ are the clusters? Although some samples are clustered together, not all show the same amount of cohesion or tightness, as defined by the amount of similarity of an object/sample within its own cluster and measured by silhouette widths. Samples with a low degree of cohesion or tightness are typically filtered out and the remaining samples define a ‘core’ sample set. Of note, we refer to a defined ‘topmost variable’ gene set as a core gene set as they are similar in their use for downstream analysis of core samples. Lastly, despite the popularity of heatmaps, information required to replicate results remains poorly documented and for this, we have implemented in NOJAH an output workflow that contains the minimum information required to replicate heatmap results.

The ‘Genome-Wide Heatmap’ analysis tab in NOJAH requires genome-wide data as input with columns representing the samples and rows, the genes. Additional details on the file format and settings are available on the NOJAH homepage. The results computed from each analysis case use are carried over as input into the next use case as part of the GWH workflow. A ‘Run Analysis’ button is available within each tab that allows the user to initiate the analysis. This feature offers the user flexibility to create multiple input parameter updates before re-running an individual analysis. Additionally, while developed as a comprehensive workflow for genome-wide heatmap analysis, each section of the workflow in Fig 1 may also be independently run.

#### Application: TCGA BRCA Expression data

Using the genome-wide TCGA BRCA RNA-Seq derived gene expression data from the 25 breast patients whose survival times we categorized into early (n= 17) and late (n= 8), we illustrate in Fig 2 the GWH workflow implemented in NOJAH. Shown in Fig 2A, is the column dendrogram based on 50,248 genes expression that shows in general a separation of samples into two clusters. While one cluster contains most samples with early survival times, the other is defined by a mixture of the two groups. Based on boxplots for the three measures of spread available in NOJAH (Fig 2B), the IQR (inter-quantile range) shows a larger spread as compared to the MAD, while the VAR approach shows the largest number of gene outliers. Therefore, we opted to use the IQR to define a topmost variable (a.k.a., ‘core’) gene set by extracting genes with an IQR above the 99^th^ percentile, resulting in 605 core genes. While a heatmap of this core gene set shows a clearer separation of samples into two clusters as compared to the genome-wide dendrogram (Fig 2A) a mixture of samples with early and late survival times remains in one cluster. As shown in Fig 2D, the consensus clustering results confirms the observed separation of samples into two clusters. Silhouette plots show samples with low widths as compared to most other samples within each cluster that suggests their removal, resulting in a defined set of 17 core samples. Using the 605 core genes with the 17 core samples, an updated heatmap shows a clear separation of samples into two clusters and further, that the clusters mostly correspond to sample groups defined by early and late survival times.

**Fig 2.**
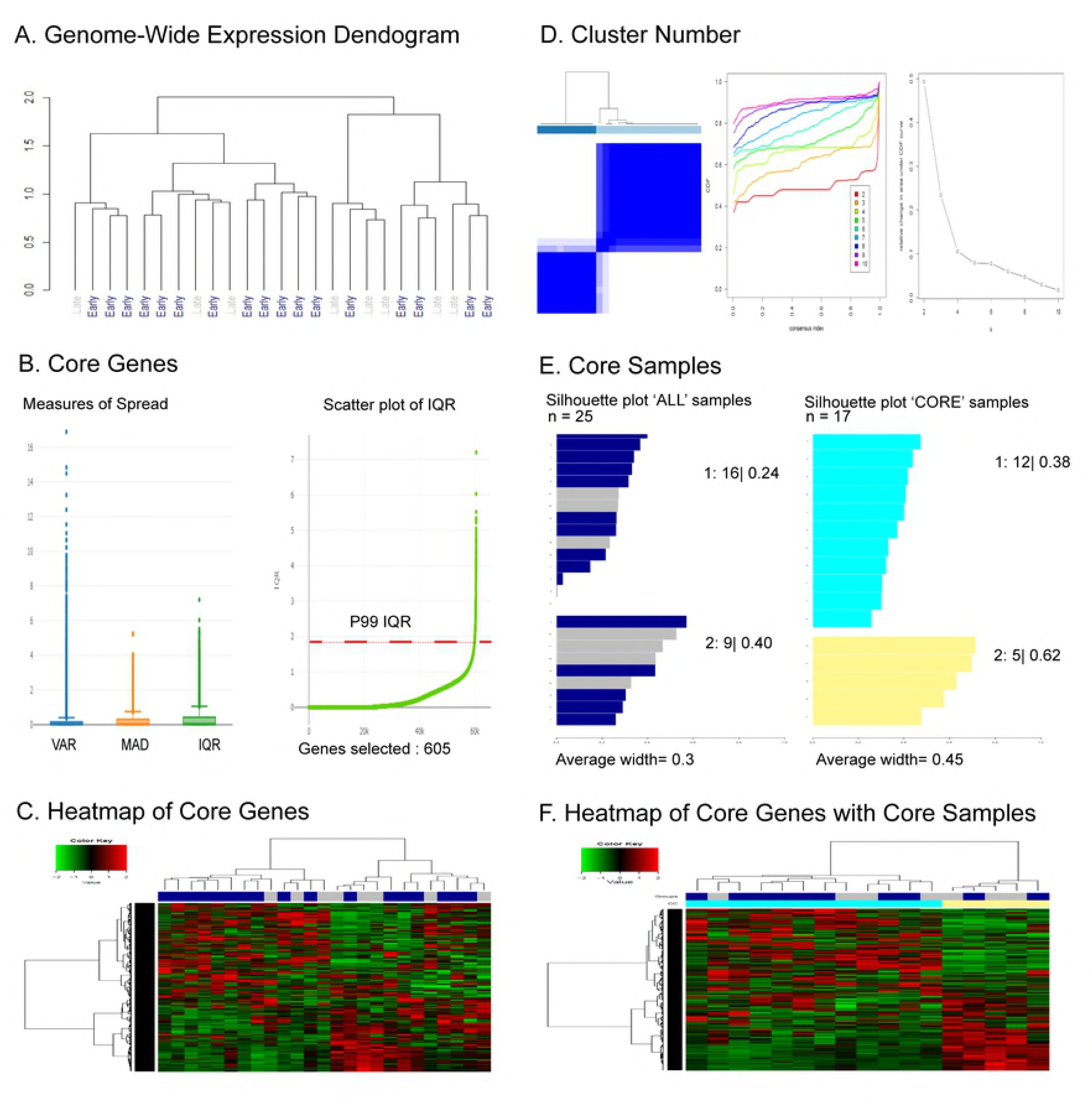
NOJAH genome-wide heatmap (GWH) Analysis of RNA-Seq derived gene expression data using TCGA BRCA expression dataset. A) Genome-wide dendrogram based on 50,248 genes using row normalization, 1-pearson correlation distance and ward.D clustering showing two clusters; one cluster containing many early survival time patients and the other, a mixture of early and late. B) *Defining Core Genes*. Distributions of measures of spread, VAR (variance), MAD (median absolute deviation), and IQR (inter-quartile range) shows IQR with fewer gene outliers as compared to VAR and a greater spread as compared to MAD. An ordered plot of IQR values for each gene shows 99^th^ percentile as a cut point to define 605 topmost variable genes (a.k.a., ‘core gene set’). C) *Heatmap of Core Genes*. Heatmap using core gene set with options: z-score based row and column normalization, 1-pearson correlation distance, and agglomerative ward.D linkage shows two distinct gene and sample clusters. D) *Defining Number of Clusters*. Results from consensus clustering using 1-Pearson correlation distance, average clustering, 80% item resampling, 100% gene samples, and agglomerative hierarchical clustering shows two clusters in the data. E) *Defining Core Samples*. Silhouette plots of samples within each of two clusters. Samples with a silhouette-width less than 0.15 and 0.34 in clusters 1 and 2 respectively were removed to define the ‘Core subset’ of 12 and 5 samples respectively. F) *Heatmap of Core Genes with Core Samples*. Updated heatmap with options: row and column z-score normalization, 1-pearson correlation distance, and agglomerative ward.D linkage clustering, based on core genes with core samples shows two distinct gene and sample cluster with most early survival time patients in one cluster and those with late times in the other.

NOJAH provides the user as output, the options selected in each step of the GWH analysis workflow to produce results, as illustrated in Fig 3 for this example. Additionally, the time elapsed in computing each step is also shown. In this case, our GWH analysis workflow runtime in total was less than two minutes. Although used for illustration, NOJAH’s GWH analysis is not limited to gene expression data but may also be applied to any genome-wide data type such as methylation, copy number, and variant, as illustrated in the combined results cluster analysis case use (Fig 4) Considering the large size of such data, NOJAH supports upload in an RDS format.

**Fig 3.**
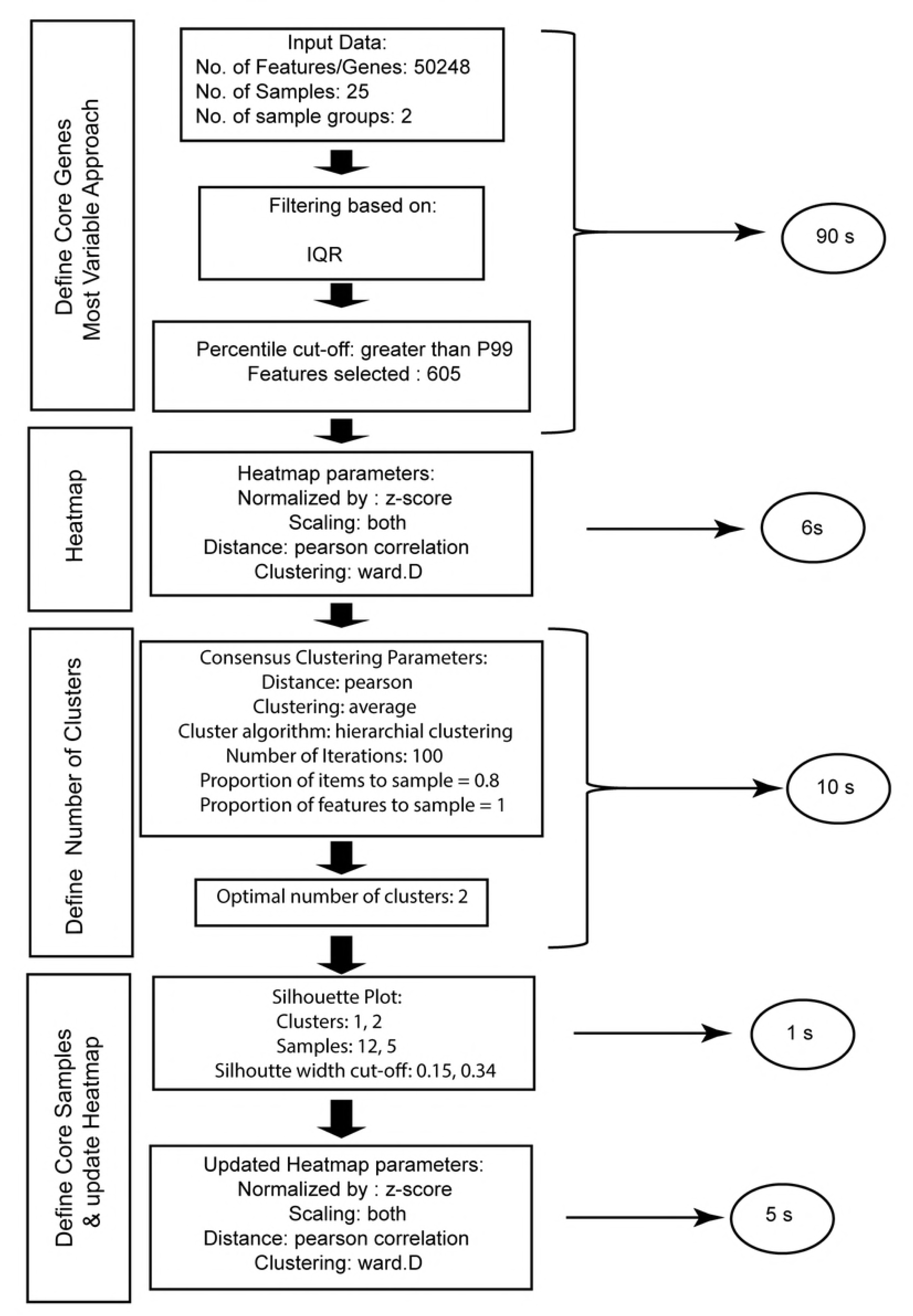
NOJAH genome-wide Heatmap (GWH) analysis output workflow using TCGA BRCA expression dataset. The optional parameters used to generate each analysis case in the GWH analysis workflow are defined as part of the output. The time in seconds (s) to run each analysis case is shown in a circle. The total time elapsed to perform a GWH analysis of RNA-Seq derived gene expression data using our example of 25 breast cancer patients was less than 2 minutes.

**Fig 4.**
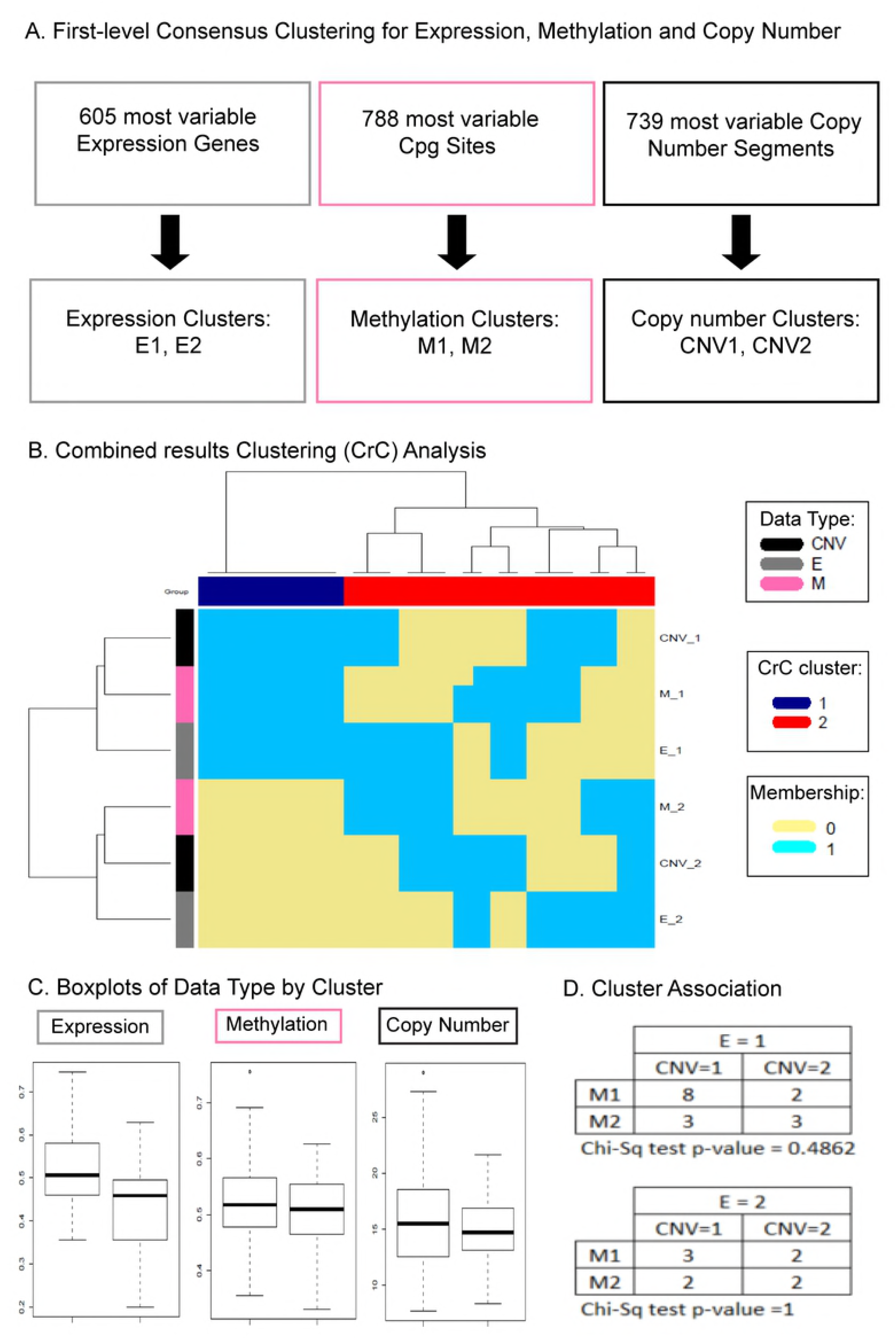
NOJAH combined results clustering (CrC) analysis of gene expression, methylation and copy number data using TCGA BRCA dataset. A) *Genome-wide Heatmap (GWH) Analysis.* A GWH analysis workflow was applied to each data type, resulting in two sample clusters based on defined most variable genes, CpG sites and copy number segments. Consensus clustering was carried out using 1-pearson correlation distance and average clustering for expression and methylation data, and canberra distance with mcquitty clustering for copy number data. For each data type, 80% sample resampling, 100% gene resampling with 100 iterations and agglomerative hierarchical clustering was performed. B) *Heatmap of cluster results*. Using a binary (0-1) matrix to indicate sample cluster membership based on individual data types, a heatmap shows two sample clusters, as also indicated by consensus clustering using the same parameters as in A, except euclidean distance and ward.D hierarchical clustering. One cluster (CrC 1 in blue) includes samples from E1, M1, and CNV1 clusters. The second cluster (CrC 2 in red) includes a mixture of samples from the various clusters. C) *Cluster Interpretation*. Boxplots of data type clusters indicated that CrC 1 includes samples with increased gene expression (E1), increased methylation (M1) and increased copy number (CNV1). A mixture of samples defined the CrC 2 cluster, including those with decreased gene expression (E2), decreased methylation (M2) and decreased copy number (CNV2). A contingency table shows many samples (n=8) with increased gene expression, increased methylation, and increased copy number.

### Combined results Cluster (CrC) analysis

A combined results cluster (CrC) analysis workflow is implemented in NOJAH based on a cluster of clusters approach [3]. Using data from several diverse genome-wide data types (e.g., gene expression, copy number, methylation, variant allele frequencies) on the same samples, a cluster of clusters approach combines separate cluster results from each platform. In specific, a binary (0-1) matrix is defined with each column denoting a sample and each row, a sub-cluster from each platform. A cluster analysis is then performed on this binary matrix of results defining a combined results cluster analysis. NOJAH’s CrC workflow also provides boxplots of sample clusters within each data type to support the user with the interpretation of combined clusters. Additionally, a contingency table analysis is also available as an option to conduct tests of association among clusters, in addition to providing the frequency distribution of samples among them.

#### Application: TCGA BRCA Expression, Methylation and Copy Number data

A GWH analysis workflow was applied separately to each of RNA-Seq derived gene expression, methylation and copy number data on the 25 breast cancer samples. Within each data type, a core gene set was defined (Fig 4A). For gene expression, IQR was used to define 605 core genes, while for copy number VAR was used to define 739 most variable, core copy number segments. In the case of methylation data, an integrated most variable analysis approach was invoked in NOJAH using a combination of both IQR and MAD measures of spread and cut off corresponding to the 99^th^ percentile applied to the combined sum of ranks of these two measures to obtain 788 most variable, core CpG sites (see supplement for details on an integrated most variable analysis approach).

Using these core sets, consensus clustering was performed on each data type to define number of clusters, resulting in k = 2 clusters in each of expression, methylation and copy number. The resulting clusters within each data type were combined into a binary matrix and a cluster analysis performed on it. Based on the CrC heatmap (Fig 4B), sample cluster 1 (CrC 1 in blue) includes samples from E1, M1 and CNV1 clusters, while cluster 2 (CrC 2 in red) includes samples from a mixture of data type clusters. **Error! Reference source not found.**The boxplots of sample groups by data type (Fig 4C) provides an interpretation for the combined clusters such that CrC 1 includes samples with increased gene expression (E1), increased methylation (M1), and increased copy number (CNV1). In contrast, CrC 2 is defined by a mixture of samples, including those with decreased gene expression (E2), decreased methylation (M2), and decreased copy number (CNV2). A contingency table, when stratified by expression clusters, shows that E1 (increased gene expression) is mostly defined by samples that also fall into both CNV1 (increased copy number) and M1 (increased methylation) clusters, whereas samples in the E2 (decreased expression) cluster are characterized by a mixture of the methylation and copy number clusters.

### Significance of Clusters (SoC) Analysis

It is often of interest to examine the statistical significance of a gene set in separating samples into groups as compared to random genes set of the same size. The significance of cluster analysis workflow uses a bootstrap approach that requires as input a user-provided genome-wide dataset with the same samples and their respective sample groupings as in the gene set of interest to test (see *supplementary methods* for details). While we illustrate gene set significance testing in NOJAH with two sample groups, the workflow may be applied to greater than two groups. The output from the SoC workflow is a table with the results from the significance testing, in addition to the heatmap of gene set of interest. Specifically, within NOJAH’s SoC workflow, a user has the option to generate a heatmap interactively using the same parameters available as in the Heatmap tab located under the GWH analysis workflow.

#### Application: TCGA BRCA Expression data

Using the 605 core genes and 17 core samples defined from the GWH analysis of the 25 TCGA BRCA expression samples dataset in Fig 1F, two sample groups were defined. By applying the SoC workflow, the 605 core genes can separate the samples into two groups, outperforming 1,000 random, 605 gene sets in this regard.

### Application

Our NOJAH application tool provides a comprehensive resource to users for conducting a genome-wide heatmap analysis. NOJAH is flexible in terms of data input, and can be applied to any data type and platform, such as mRNA expression, miRNA expression, methylation, copy number or variants. Additional features in NOJAH include interactive settings for defining core genes and core samples and combined results clustering, along with the flexibility to include phenotype information through use of a color bar. While we have demonstrated the utility of NOJAH using a TCGA BRCA data set of gene expression, any high-dimensional quantitative data may be used as input.

## Discussion

Identification of gene signatures is crucial in cancer genomics. Prognostic gene signatures within a cancer type constitute a set of genes whose expression changes reveal important information about tumor diagnosis, prognosis and even therapeutic response [9, 12]. The dependence on the use of heatmaps to apply published gene signatures for tumor subtyping is increasing and along with it, the challenges in obtaining results. With a comprehensive workflow in hand as a single application tool, there is little room for computational error by invoking several separate tools to accomplish the end task of applying a gene signature for tumor subtyping. Additionally, with a workflow that includes as output the parameters used to obtain results, the replicability of them is more feasible than with documenting several steps from several programs and approaches.

While there exists many tools for heatmap construction, each of them has certain limitations. As example, a recently developed heatmap tool, shinyHeatmap [13], was unable to compute results for our CoMMpass RNA-Seq gene-level expression genome-wide dataset of 560 samples with 60,000 rows. Another commonly used heatmap tool, Morpheus (https://software.broadinstitute.org/morpheus/), allows the user to filter genes based on numerous descriptive measures (e.g., VAR, MAD, maximum, minimum, mean) without providing much evidence as to which may be more appropriate for a dataset. In our NOJAH tool, the user can access the data distribution based on the interactive boxplots and scatterplots to make a more informed choice about which filters and associated percentile cut-off may be more appropriate for their data, and includes the option for a combination of methods which is especially helpful in the case of sparse data. Neither heatmap tool has a built-in functionality to perform a systematic, comprehensive cluster analysis. Each tool is based on a single hierarchical clustering or k-means clustering approach and does not enable further examination of either the number of clusters or the tightness of clusters. NOJAH is a heatmap analysis tool enabling the user to implement a comprehensive workflow in which a user can not only perform hierarchical clustering but also confirm the number of clusters, define a core sample set to improve the tightness of clusters and re-create heatmaps, all using an interactive, point and click functionality.

Our NOJAH application tool provides researchers with a comprehensive workflow in which to apply known and identify novel gene signatures, and equips them with an easy access tool to perform additional timely analysis, such as in combining cluster results from multiple genomic platforms. In NOJAH we provide descriptives that enable the user to interpret the clusters within each data type in order to provide an overall interpreation for the cluster of clusters obtained, something that was absent in the literature. For example, in the four-data type (methylation, mRNA, copy number, microRNA) cluster of clusters analysis performed on lower-grade gliomas (LGG) reported by The Cancer Genome Atlas Research Network, three resulting clusters were reported [3]. Although a strong correlation between wild-type IDH (no mutation in IDH1 or IDH2) and IDH-1p/19q codeletion was observed as associated with one of the clusters, there was minimal to no information about how to interpret the actual resulting clusters in terms of direction of change as either increased/decreased among the data types. In addition to providing such information, NOJAH also includes a contingency table analysis for tests of association among clusters.

Some available R packages such as heatmap3 allow the user to test the statistical significance of the annotated sample groups with the clusters based on a chi-squared test. The SoC workflow inn NOJAH has taken this a step further by testing the statistical significance of the gene set in separating samples into groups as compared to random gene sets of the same size using a bootstrap approach.

Finally, NOJAH helps to minimize one of the most commonly observed issues in heatmap construction that no other current heatmap tool offers (e.g. Morpheus, shinyHeatmap [13], ClustVis [14], WebGimm [15]), which is the lack of sufficient information for replicating a heatmap. A pdf file with the exact settings/parameters that were used to generate the heatmap along with the row and column dendrograms are made available for download in pdf format. NOJAH also provides to the users a detailed galaxy-like workflow [16] in the GWH tab with the exact settings used in each step of the workflow.

Our NOJAH tool is the only comprehensive heatmap and cluster tool, making it a heatmap analysis application that harbors three independent workflows for genome-wide data: heatmap analysis, combined results cluster analysis, and significance of cluster analysis. The implementation of so many tools in a single interface with a workflow eliminates the requirement to install and learn how to use and output results from several codes, requiring only a stable web connection to load. NOJAH is thus truly a one-stop shop for performing a heatmap analysis. In summary, NOJAH is a freely available, both as a web interface and stand-alone version, user-friendly tool developed in response to the need for updated approaches to address genome-wide cluster analysis of single and multiple data types, and significance testing of resulting gene signatures, in a computing environment that fosters replicability and accessibility through a point and click functionality.

### Implementation

NOJAH is a web-interface developed using the Shiny R package [17] hosted on a private Centos OS server and requires only a stable internet connection to run. The source code is written in the R programming language (https://www.r-project.org/) and is freely available to download from the GitHub (https://github.com/bbisr-shinyapps/NOJAH/). The main R packages used in NOJAH include: heatmap.2, gplots, ConsensusClusterPlus [2], dendextend [18], and plotly [19]. NOJAH was tested using google chrome and firefox on a 64-bit, x64-based processor Windows 10 Enterprise machine with 32GB of RAM and an Intel(R) Core(TM) i7-7820HQ CPU at 2.90 GHz.

### Limitations

The tool is hosted on a virtual private server to maintain the web application for the next five years which we intend to upgrade based on the inbound load. The speed at which the tool generates results will depend upon the user’s internet speed and the size of the input data. It may take a few seconds to a few minutes for the analysis to finish running in the background and display the results, depending on the size of the input data. For example, when generating a heatmap for data with 1000-1500 genes, the output is displayed within seconds, but when performing a genome-wide heatmap analysis using datasets that contain between 20,000-60,000 genes, it may take about two minutes to generate the results. Large datasets (>50 MB) may be converted to RDS format before upload to overcome size issues but for very large datasets that exceed the shiny upload limit (>500MB), a GitHub version with the R source code is made available to run on the user’s local computer using the R command line, though the speed will depend on the user’s local computer processing speed.

NOJAH requires the input data to be placed in a specified format that is documented on the tool’s homepage. When uploading your own data, the user must change the defaults to reflect the row and column numbers where the numeric data starts. For example, one dataset may have information on the status of estrogen receptor (ER), progesterone receptor (PR), and human epidermal growth factor receptor type 2 (HER2), which would be represented on the top of the heatmap just below the column dendrogram and a column with the gene groups information in addition to the gene names. In such cases, the numeric data starts at row number five and column number three. In another example, the user may not have any additional sample information to display but the same gene group information, thus the numeric data may start at row number three and column number three.

In the current version of NOJAH, a maximum of 10 sample groups and six feature groups may be compared. The color of the groups to be displayed on the heatmap are fixed. We intended to increase the number of feature and sample groups as well the choice of the color for the groups as future options. Additionally, at present, a maximum of three data types can be used in the ‘CrC’ workflow and we are working to extend this option to allow a greater number. Also, in a future upgraded version of NOJAH, we intend to allow the user to remove any outlier sample(s) from each CrC tab that may be indicated by the silhouette plots to perform a more comprehensive CrC analysis.

## Acknowledgments

We would like to thank the Cancer Informatics Core of the Winship Cancer Institute of Emory University for hosting NOJAH on the CentOS Virtual Machine. We would like to extend a special thanks to Kenneth Buck for his assistance with building and configuring the server.

## Funding

Research reported in this publication was supported in part by a Winship Glenn Family Breast Cancer award (Kowalski) and the Biostatistics and Bioinformatics Shared resource of Winship Cancer Institute of Emory University and National Institutes of Health /National Cancer Institute under award number P30CA138292.

## Conflict of Interest

none declared.

## Author Contributions

Conceptualization: JK

Data Curation: MR, BD

Formal Analysis: MR, BD, JK

Methodology: JK

Software: MR

Supervision: JK

Validation: MR, BD, JK

Visualization: MR, BD, JK

Writing – Original Draft Preparation: MR

Writing – Review & Editing: BD, JK

